# Tuning epithelial cell-cell adhesion and collective dynamics with functional DNA-E-cadherin hybrid linkers

**DOI:** 10.1101/2021.09.27.462021

**Authors:** Andreas Schoenit, Cristina Lo Giudice, Nina Hahnen, Dirk Ollech, Kevin Jahnke, Kerstin Göpfrich, Elisabetta Ada Cavalcanti-Adam

## Abstract

The binding strength between epithelial cells is crucial for tissue integrity, signal transduction and collective cell dynamics. However, there is no experimental approach to precisely modulate cell-cell adhesion strength at the cellular and molecular level. Here, we establish DNA nanotechnology as tool to control cell-cell adhesion of epithelial cells. We designed a DNA-E-cadherin hybrid system consisting of complementary DNA strands covalently bound to a truncated E-cadherin with a modified extracellular domain. DNA sequence design allows to tune the DNA-E-cadherin hybrid molecular binding strength, while retaining its cytosolic interactions and downstream signaling capabilities. The DNA-E-cadherin hybrid facilitates strong and reversible cell-cell adhesion in E-cadherin deficient cells by forming mechanotransducive adherens junctions. We assess the direct influence of cell-cell adhesion strength on intracellular signaling and collective cell dynamics. This highlights the scope of DNA nanotechnology as a precision technology to study and engineer cell collectives.

## Main

Epithelial cells are linked to one another to maintain tissue structural integrity and to respond dynamically to events which require coordinated behavior, like morphogenesis or collective migration (Friedl and Gilmour, 2009). Adherens junctions (AJs) mediate strong cell-cell adhesion and are especially important for the transduction of mechanical signals between cells, which govern collective dynamics (Halder et al., 2012; Ladoux and Mège, 2017; Lecuit and Yap, 2015; Mui et al., 2016). To establish the physical link, the adhesive receptors cadherins form *trans*-dimers with cadherins of neighboring cells. Epithelial cadherin (E-cadherin) is crucial for the epithelium integrity, and its loss is associated with different forms of cancer and the acquisition of invasive properties (Berx and van Roy, 2009). E-cadherin dimerization leads to downstream signaling, which involves the recruitment of other AJ proteins, e.g. catenins, and remodeling of the actin cytoskeleton (Harris and Tepass, 2010; Yap et al., 2015). At AJs, mechanical cues are translated into biochemical signals, which regulate fundamental cellular processes like proliferation and cell fate (Dasgupta and McCollum, 2019; Halder et al., 2012).

The investigation of the influence of cell-cell adhesion and its mechanical regulation on these processes requires control over AJ assembly and functionality, which is mainly achieved by modulating AJ protein expression levels (Gumbiner, 1996). Mutations in cadherins (Handschuh et al., 1999) or RNA interference (Das et al., 2015) provide some control over cell-cell adhesion strength, but these approaches require extensive tuning depending on the cell type and experimental conditions. The depletion of calcium ions, required for cadherin dimerization (Jain et al., 2020; Kim et al., 2011), or recently reported optochemical and optogenetic approaches (Cavanaugh et al., 2020; Kong et al., 2019; Ollech et al., 2020) facilitate the spatiotemporal control over cell-cell adhesion assembly and disassembly. However, the influence of the molecular binding strength between cells remains unknown, since there is no method to precisely control it.

DNA nanotechnology allows for the programmable generation of molecular architectures with a sequence-tunable binding strength (Pinheiro et al., 2011; Seeman and Sleiman, 2017). Due to this versatility, combined with a large toolbox of chemical functionalization options, several applications have been presented in cell biology studies (Schoenit et al., 2021). DNA is commonly anchored on the cell membrane by hydrophobic moieties, like cholesterol or fatty acids that are covalently linked to the DNA (You et al., 2017). It has been used to facilitate artificial cell-cell adhesion in non-adherent or suspended cells (Borisenko et al., 2009; Gartner and Bertozzi, 2009), even with complex DNA nanostructures like DNA origami (Akbari et al., 2017; Ge et al., 2020). Furthermore, it has been reported that the control over the binding strength can be achieved by varying the DNA concentration (Hoffecker et al., 2019). Moreover, DNA allows to program the cellular organization in bottom-up tissue assembly (Todhunter et al., 2015) or to report forces within cellular monolayers (Zhao et al., 2020). However, since DNA strands alone inserted into the cellular membrane do not interact with intracellular structures (Gartner and Bertozzi, 2009), the functionality of the artificial DNA link remains questionable. It is unclear whether downstream signaling is maintained upon artificial DNA-mediated cell-cell adhesion and its use in cell collectives remains largely unexplored.

Here, we present a functional DNA-E-cadherin hybrid to tune adhesion strength in epithelial cell collectives. By linking DNA to a truncated E-cadherin construct, we ensure a link with the intracellular machinery, while benefitting from the DNA sequence-dependent adhesion strength. Using force spectroscopy, we verify an increased cell-cell adhesion strength upon DNA linker addition, which can be reversed by using DNA strand displacement reactions. Furthermore, we demonstrate the recruitment of AJ proteins, which eventually leads to mechanosensing. Finally, we apply the DNA-E-cadherin hybrid to investigate the influence of cell-cell adhesion strength on collective dynamics in migrating epithelial monolayers.

First, we set out to achieve a precisely controllable semi-synthetic, yet functional cell-cell adhesion linker for epithelial cells. While DNA-mediated cell-cell adhesion can be easily facilitated by a cholesterol-tagged DNA linker in non-adherent cells (**Figure S1**), this approach is not suitable for functional studies that require downstream signaling, like for epithelial cell collectives. We thus designed a DNA-E-cadherin hybrid system. Our approach combines the tunability and versatility of DNA with the intracellular signaling capabilities of the adhesive receptor E-cadherin, which binds *via* the AJ proteins catenins to the actin cytoskeleton (**Figure 1 A**). *Trans-* and *cis-*clustering with other E-cadherins is facilitated by the two outer extracellular domains EC1 and EC2 (Boggon et al., 2002). Additionally, adhesion-independent E-cadherin clustering can be promoted, to a limited extent, by the actin cytoskeleton (Wu et al., 2015). To achieve control over the extracellular interactions involved in cell-cell adhesion, we replaced these domains with a SNAP-tag inserted into the sequence between R154 and N376, thereby maintaining the sequence for correct extracellular expression and further protein modification. The SNAP-tag allows fast and highly specific binding of any molecule functionalized with its ligand benzylguanine (Keppler et al., 2003). The expression of SNAP-E-cadherin in A431D cells, which lack endogenous expression of E-cadherin (Lewis et al., 1997), was successful but did not facilitate cell-cell adhesion (**Figure S2 A**). We designed a 45 base pair long DNA linker consisting of identical benzylguanine-tagged anchor strands, which covalently bind to the SNAP-tag and hybridize with two complementary linker strands. The sequence-complementary part exhibits a calculated molecular binding strength of 17.7 kcal/mol (Zadeh et al., 2011) and facilitates *trans-*dimerization of SNAP-E-cadherin. Furthermore, the DNA was functionalized with a fluorophore (e.g. Cy5) for visualization (**Figure 1 A**).

**Figure 1:**
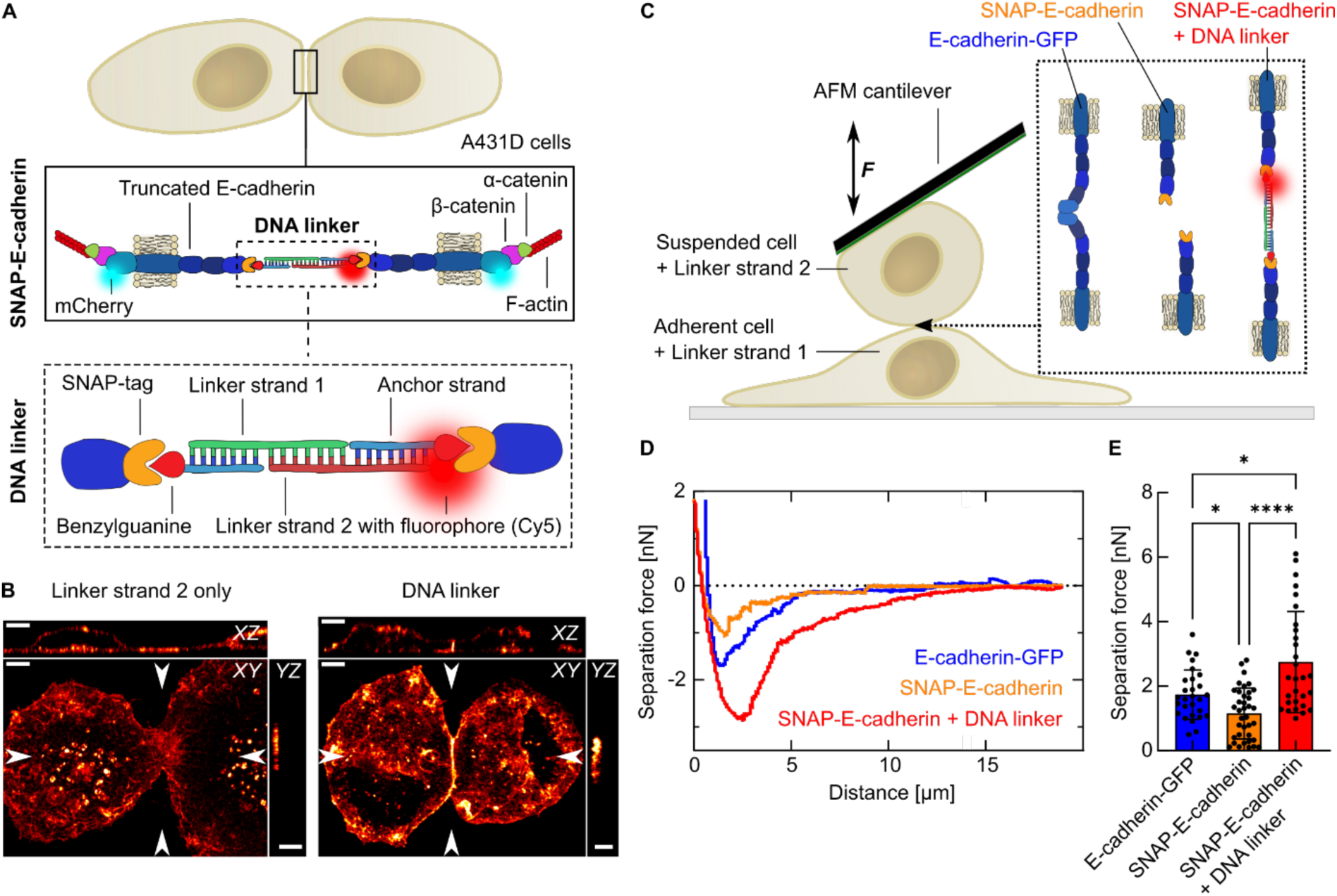
Cell-cell adhesion is facilitated by a DNA-E-cadherin hybrid linker. **A** Sketch of the DNA-E-cadherin hybrid linker system. Epithelial A431D cells express a truncated E-cadherin, where the extracellular domains EC1 and EC2 are replaced by a SNAP-tag. The intracellular domain is labelled with mCherry and binds via the proteins β-catenin and α-catenin to the F-actin cytoskeleton. SNAP-E-cadherins from neighboring cells form *trans-*dimers in presence of the DNA linker. The SNAP-tag allows the binding of a 15 bp long anchoring DNA strand functionalized with benzylguanine. Two complementary 30 bp long linker strands are bound to the anchoring strand to form a duplex. The linker strands can be tagged with a fluorophore (e.g. Cy5) for visualization. **B** Whole-cell 3D reconstructions of A431D cells expressing SNAP-E-cadherin. Cells were incubated with only linker strand 2 carrying Cy5 (*left*) or the complete Cy5-tagged DNA linker (*right*). Maximum projection and orthogonal slices through the positions indicated by arrows are shown for the Cy5 channel. Scale bars, 5 µm. **C** Single cell force spectroscopy using AFM. Sketch of the experimental setup: Adherent cells are pre-incubated with Linker strand 1 for 1h. Suspended cells pre-incubated with Linker strand 2 are captured with the AFM cantilever. The cells are brought in contact and the separation forces are measured. **D** Representative separation force curves for cells expressing full-length E-cadherin-GFP (*blue*), SNAP-E-cadherin (*orange*) and SNAP-E-cadherin incubated with the DNA linker (*red*). Cell contact duration is 5 s. The separation force is the minimum of the curve. **E** Comparison of the separation forces. Bars show the mean value. Error bars show the standard deviation. Plots generated from *N* = 3 independent experiments. Number of measured cells: *n*(E-cadherin-GFP) = 28. *n*(SNAP-E-cadherin) = 37. *n*(SNAP-E-cadherin + DNA linker) = 29. (*) p-value between 0.1 and 0.01. (****) p-value < 0.0001. Multiple ANOVA tests with Welch’s correction. Alpha was set to 0.05.

The addition of only Linker strand 2 did not induce cell-cell contacts as no DNA duplex between two cells can be formed (**Figure 1 B, Figure S2 A, Supporting Video S1**). In contrast, the addition of the complete DNA linker led to an accumulation of both DNA (**Figure 1 B**) and SNAP-E-cadherin construct (**Figure S2 A, Video 1**) at the cell-cell interface. Furthermore, we observed the formation of a straight cell-cell junction with an increased height compared to the non-functional single linker strand. The DNA linker was stable at the cellular membrane for several hours, but its presence decreased over time, likely due to internalization of the DNA-E-cadherin hybrid receptors (**Figure S2 B, C**). To demonstrate that DNA-mediated cell-cell adhesion on the molecular level translates to an increased cell-cell adhesion strength on the cellular level, we performed single cell force spectroscopy measurements using an atomic force microscope (AFM). For this purpose, we probed the separation forces between a suspended cell bound to the AFM cantilever and an adherent cell (Friedrichs et al., 2013; Panorchan et al., 2006) (**Figure 1 C, Figure S3 A**). We compared A431D cells that either expressed full-length E-cadherin-GFP or the truncated SNAP-E-cadherin construct. In the latter case, adherent cells and suspended cells were separately pre-incubated with one of the complementary DNA linker strands, respectively. The captured cell was pushed on the adherent cell to probe cell-cell interactions and the separation force was measured. The force spectroscopy measurements revealed that the truncated SNAP-E-cadherin, in the absence of the DNA linker, results in significantly weaker cell-cell adhesion than the E-cadherin-GFP (1.7 ± 0.8 nN versus 1.2 ± 0.8 nN, respectively). These values are in the same range as reported for other cadherin-dependent single cell force spectroscopy experiments in different cell types (Fichtner et al., 2014; Pawlizak et al., 2015). Importantly, the addition of the DNA linker to SNAP-E-cadherin-expressing cells, and therefore the assembly of the DNA-E-cadherin hybrid system, led to a significantly increased adhesion strength of 2.8 ± 1.6 nN (**Figure 1 D, E**). We observed this trend for different contact times (2, 5 and 10 seconds, **Figure 1 E, Figure S3 B, C**), which demonstrates that the DNA linker leads to fast and strong cell-cell adhesion.

Besides the controlled inducibility, another advantage of using a DNA linker is the reversibility of the linkage, i.e. the DNA duplex can be opened by toehold-mediated strand displacement (Zhang and Winfree, 2009). To enable strand displacement, we adapted the linker design by adding a 15 nucleotide long sequence overhang to Linker strand 2. This single-stranded DNA toehold-overhang cannot hybridize with Linker strand 1 since it is not complementary. Additionally, we designed an invader strand, which is complementary to Linker strand 2 including the toehold sequence. Its binding affinity to the toehold-modified Linker strand 2 is thus higher than the affinity between the two linker strands (**Figure 2 A**). Added in excess, the invader strand rapidly removed the toehold-modified linker strand within one minute to open the DNA linker. The toehold-modified linker strand then diffuses out of the focal plane resulting in a increased background together with internalized strands (**Figure 2 B, Video 2**). Hence, our approach provides temporal control to reversibly turn cell-cell adhesion on and off.

**Figure 2:**
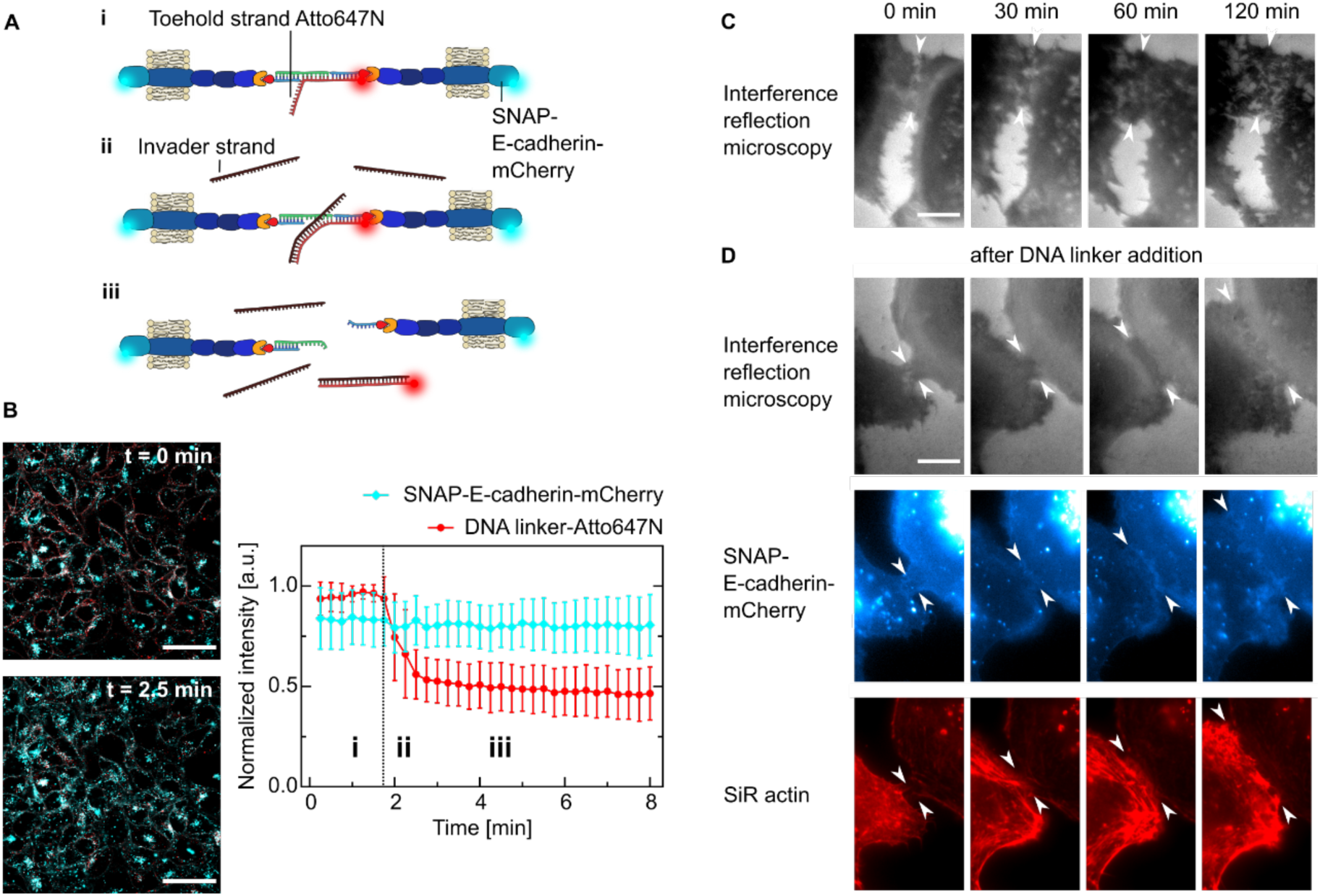
Dynamic control of cell-cell adhesion. **A** Sketch showing the reversibility of the cell-cell linkage by using toehold-mediated strand-displacement. A431D cells are linked with a DNA strand which contains a free overhang (toehold) functionalized with Atto647N (**i**). Upon addition of an invader strand in excess (**ii**), the linker strand is displaced and the link between the cells is broken (**iii**). **B** Representative confocal images of cells expressing SNAP-E-cadherin-mCherry (*cyan*) incubated with the DNA linker-Atto647N (*red*) before and after the addition of the invader strand. Scale bar, 50 µm. Quantification of the fluorescence intensity normalized to the maximum value over time of SNAP-E-cadherin-mCherry and DNA linker-Atto647N. The invader strand is added at t = 1.8 min, indicated by the dotted line. Mean values are plotted and error bars indicate the standard deviation. Plots generated from *N = 3* independent experiments and *n* = 31 measurements. **C** Live-cell time-lapse snapshots of A431D cells expressing SNAP-E-cadherin-mCherry imaged by interference reflection microscopy without addition of the DNA linker. Images representative of *N = 2* independent experiments. Scale bar, 10 µm. **D** Live-cell time-lapse snapshots showing the formation of a cell-cell junction between A431D cells expressing SNAP-E-cadherin-mCherry after the addition of the DNA linker. Top row, interference reflection microscopy; middle row, SNAP E-cadherin (*blue*); bottom row, actin cytoskeleton labeled with SiR actin (*red*). Images representative of *N = 2* independent experiments. Scale bar, 10 µm.

Having demonstrated that the DNA-E-cadherin hybrid allows reversible cell-cell adhesion, we next determined whether the linkage between cells is functional and enables downstream signaling. Unlike commonly used strategies where cells were linked using sDNA functionalized with e.g. hydrophobic tags but without a suitable transmembrane domain (Akbari et al., 2017; Borisenko et al., 2009; Gartner and Bertozzi, 2009; Hoffecker et al., 2019; Todhunter et al., 2015), we particularly designed the DNA-E-cadherin hybrid system to preserve the transmembrane and the intracellular cadherin domains which mediates the connection to the cytoskeleton. We therefore assessed the cellular reaction to the DNA linker addition qualitatively by live-cell microscopy. While the contact length between neighboring cells in absence of the DNA linker did not change over time (**Figure 2 C**), we observed a three-fold increase of the contact length within the first hour after DNA linker addition. This was accompanied by an accumulation of SNAP-E-cadherin at the interface of the cells as well as actin remodeling, which points to an intracellular response to the extracellular signal (**Figure 2 D, Video 3**).

Since the use of the DNA-protein hybrid enabled us to investigate, for the first time, the intracellular response to a DNA-based cell-cell linker, we visualized the subcellular localization of E-cadherin using stimulated-emission-depletion (STED) microscopy. Clustering of E-cadherin-GFP led to the formation of a pronounced and straight AJ (**Figure S4 A**). No defined junction was detectable for cells expressing the non-functional SNAP-E-cadherin (**Figure S4 B**). The addition of the DNA linker resulted in an accumulation of SNAP-E-cadherin at the cell-cell interface, resembling E-cadherin-GFP but with smaller and less organized clusters, reflecting the absence of the EC1 and EC2 protein domains and the contribution of the actin cytoskeleton (**Figure S4 C**). Beyond receptor clustering, assembly of functional AJs is characterized by the presence of E-cadherin/β-catenin complexes, which establish a link via α-catenin to the actin cytoskeleton (Harris and Tepass, 2010). Here, the E-cadherin/β-catenin complex is crucial for the mechanical stability of AJs (Bertocchi et al., 2017; Orsulic et al., 1999) (**Figure 3 A**). Successful formation of AJs leads to actin remodeling, where the actin fibers are aligned parallel to the junction to stabilize it (Escobar et al., 2015; Harris and Tepass, 2010; Mège et al., 2006) (**Figure 3 B**). For the bare SNAP-E-cadherin, we could neither detect co-clustering with β-catenin (**Figure 3 C**) nor the parallel alignment of actin fibers at the cell-cell junction (**Figure 3 D**). Thus, SNAP-E-cadherin by itself was not functional and did not facilitate AJ downstream signaling. In contrast, addition of the DNA linker triggered AJ-dependent signaling. Co-clustering with β-catenin (**Figure 3 E**) as well as the actin distribution at the cell-cell junction (**Figure 3 F**) were similar to the full-length E-cadherin, thus showing functional linkage between cells.

**Figure 3:**
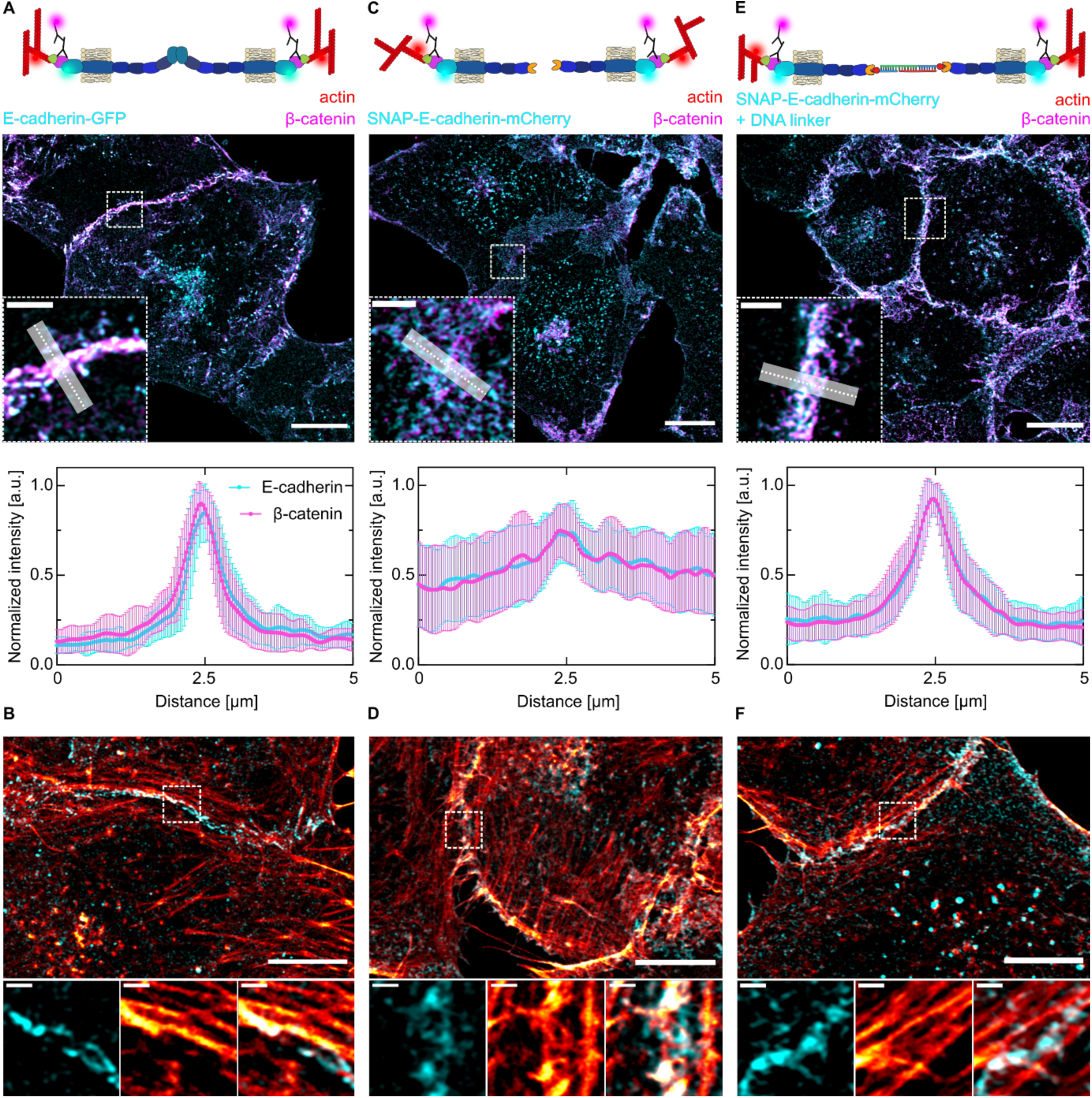
Downstream signaling of SNAP-E-cadherin after DNA linker addition. **A, C, E** Sketches of the different cell-cell adhesions between A431D cells expressing full-length E-cadherin-GFP (**A**) or SNAP-E-cadherin-mCherry without (**C**) or with the DNA linker (**E**). Airyscan confocal images of the subcellular localization of E-cadherin (*cyan*) and β-catenin (*magenta*) visualized by indirect immunostaining. Scale bars, 10 µm. Zoom-ins at the area indicated by the dashed line. Scale bars, 2 µm. Quantification of the fluorescence distribution normalized to the maximum intensity at the cell-cell interface generated by averaging the fluorescence intensity of multiple line plots (length = 5 µm, width = 1 µm). Error bars correspond to the standard deviation. *N* = 3 independent experiments and *n*(E-cadherin-GFP) = 20, *n*(SNAP-E-cadherin-mCherry) = 22 and *n*(SNAP-E-cadherin-mCherry + DNA linker) = 23 measurements. **B, D, F** Airyscan confocal images of the actin cytoskeleton (*red*) of A431D cells expressing full-length E-cadherin-GFP (**B**) or SNAP-E-cadherin-mCherry without (**D**) or with the DNA linker (**F**). Scale bars, 10 µm. Zoom-ins at the area indicated by the dashed line. Scale bars, 1 µm.

Mechanical signals are key for the determination of cell fate. An important function of AJs is sensing mechanical signals and transducing them from the outside to the inside of the cell via translation into biochemical signals (Charras and Yap, 2018). We thus investigated if our DNA-E-cadherin hybrid system is capable of mechanotransduction of intracellular downstream signaling. It is known that mechanotransduction involves the activity of the transcription regulator Yes-associated protein (YAP) and influences its subcellular localization (Halder et al., 2012). YAP is excluded from the nucleus and translocated to the cytosol upon phosphorylation, which is induced through mechanical cues from surrounding cells, sensed at functional and mechanically active AJs (Aragona et al., 2013; Benham-Pyle et al., 2015). We thus quantified YAP distribution for DNA-linked cells in comparison to the controls. In contrast to cells expressing E-cadherin-GFP, we observed the nuclear accumulation of YAP in cells expressing only SNAP-E-cadherin (**Figure 4 A**). We assessed the nuclear to cytosolic ratio of YAP (**Figure 4 B**) and quantified the fractions of areas within the monolayer where YAP was predominantly localized in the nucleus, the cytosol or both (**Figure 4 C**). The nuclear to cytosolic ratio increased from 0.7 ± 0.2 for E-cadherin-GFP expressing cells to 1.1 ± 0.3 when SNAP-E-cadherin was expressed. Since a low nuclear to cytosolic ratio indicates YAP exclusion, it shows that mechanotransduction is compromised in SNAP-E-cadherin expressing cells. The nuclear exclusion of YAP from cells in which DNA-E-cadherin hybrid was assembled, demonstrated that the mechanical signal from the cell peripheries was transduced into the cytosol and to the nucleus. Using DNA linkers with different hybridization strengths (3.2, 10.4 and 17.7 kcal/mol) revealed that in cells adhering with the 10.4 kcal/mol linker (ratio: 0.7 ± 0.1) mechanotransduction was more pronounced than in cells adhering with the 3.2 kcal/mol linker (ratio: 0.9 ± 0.1). This is reflected in the level of nuclear exclusion of YAP, which is similar to the one observed in cells expressing the full-length E-cadherin (**Figure 4 B, C**). The strongest linker did not lead to a further decrease of the nuclear to cytosolic ratio (0.7 ± 0.1), indicating that cellular mechanotransduction was fully achieved with the 10.4 kcal/mol linker. Note that the only difference between the 3.2 kcal/mol and the 10.4 kcal/mol linker is an increased length of only two base pairs. Thus, the DNA-E-cadherin hybrid facilitates physiological outside-in signaling downstream of E-cadherin, which can be fine-tuned through subtle differences in the DNA hybridization strength.

**Figure 4:**
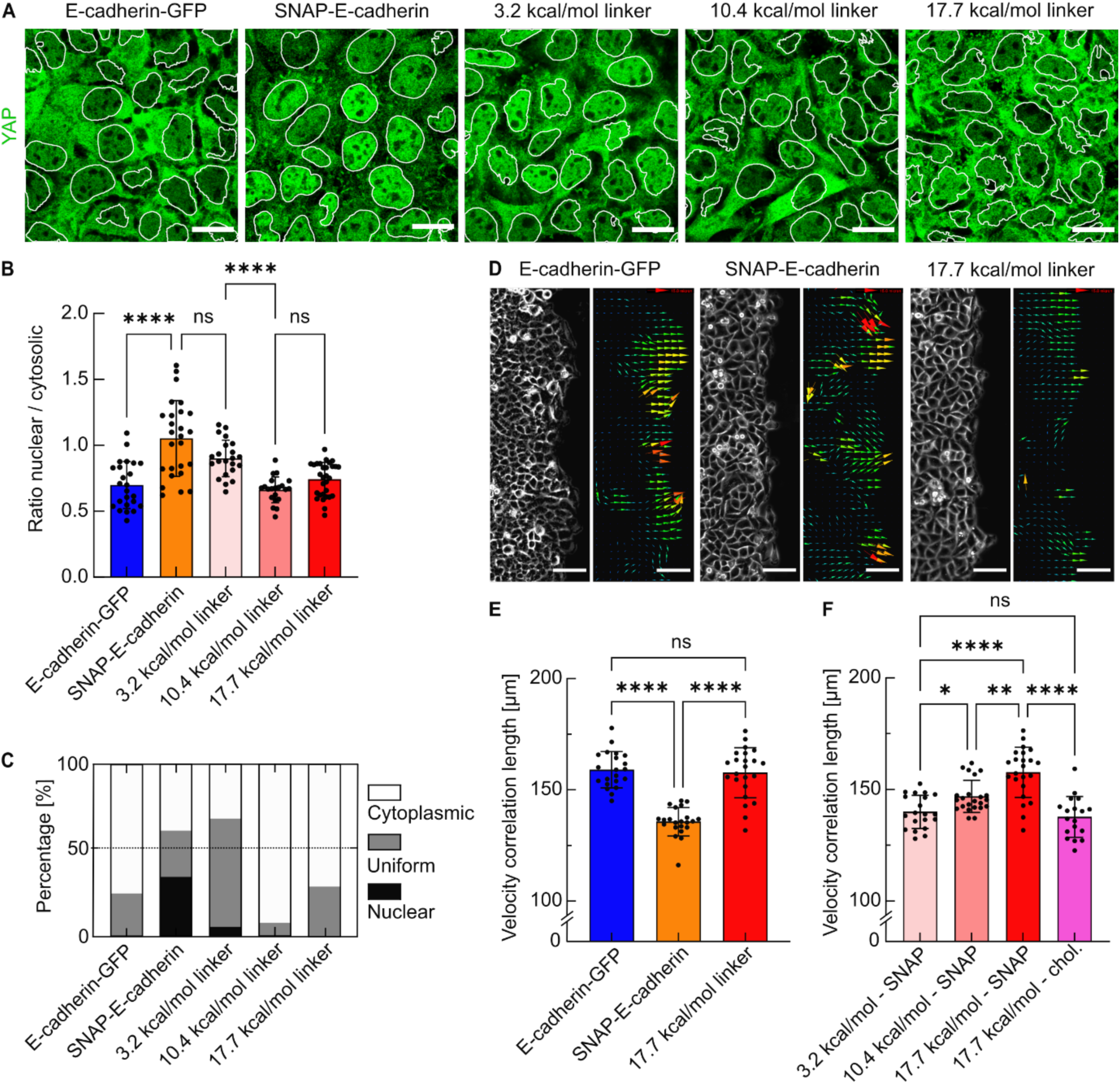
Effect of the DNA linker on cell-cell signaling and epithelial collective dynamics. **A** Confocal images showing YAP (*green*) with the nuclear segmentation (*white outlines*) for cells expressing E-cadherin-GFP or SNAP-E-cadherin after the incubation with DNA linkers of different hybridization strengths (3.2, 10.4 or 17.7 kcal/mol). Scale bars, 20 µm. **B** Quantification of the nuclear- -to-cytosolic intensity ratios *r* of YAP. Each datapoint represents one field of view containing 25 to 30 cells. Bars show the means, error bars are standard deviations. **C** Quantification of the subcellular localization of YAP classified according to the ratios *r* between nuclear and cytosolic YAP intensities. Nuclear: *r* ≥ 1.15; uniform *r* > 0.85; cytosolic: *r* ≤ 0.85. *N* ≥ 3, data derived from *n*(E-cadherin-GFP) = 24 measurements. *n*(SNAP-E-cadherin) = 25. *n*(SNAP-E-cadherin + 3.2 kcal/mol linker) = 22. *n*(SNAP-E-cadherin + 10.4 kcal/mol linker) = 21. *n*(SNAP-E-cadherin + 17.7 kcal/mol linker) = 29. **D** Brightfield and vector fields generated by particle image velocimetry (PIV) of the migration front of A431D cells expressing E-cadherin-GFP, SNAP-E-cadherin or SNAP-E-cadherin incubated with the 17.7 kcal/mol DNA linker. Scale bars, 100 µm. **E** Comparison of the velocity correlation lengths between migrating cells. Every data point shows the average correlation length of one field of view from t = 3. 5 h to t = 7.3 h after removing the confinement, corresponding to 24 timepoints. n*FOV* shows the number of analyzed fields of view from at least 3 independent experiments. n*FOV*(E-cadherin-GFP) = 21, n*FOV*(SNAP-E-cadherin) = 22, n*FOV*(SNAP-E-cadherin + DNA linker) = 23. **F** Velocity correlation length for the variation of the DNA hybridization strength (3.2, 10.4 or 17.7 kcal/mol) as well as the anchoring method (SNAP-E-cadherin or direct functionalization of the anchor strand with cholesterol). *N* ≥ 3, n*FOV*(3.2 kcal/mol - SNAP) = 19, n*FOV*(10.4 kcal/mol - SNAP) = 23, n*FOV*(17.7 kcal/mol - SNAP) = 23 (same data set as in **E** shown again for better comparability), n*FOV*(17.7 kcal/mol - chol) = 18. ns no significance. (*) p-value between 0.1 and 0.01. (**) p-value between 0.01 and 0.001. (****) p-value < 0.0001. Multiple ANOVA tests with Welch’s correction. Alpha was set to 0.05.

Since our approach allows to modulate not only cell-cell adhesion but also associated downstream signaling in cells, we next investigated how this modulated strength would impact epithelial collective dynamics on the multicellular scale. Therefore, we performed collective migration assays to assess long-range interactions between cells, which are also crucial in e.g. wound healing or morphogenesis (Ladoux and Mège, 2017; Lecuit and Yap, 2015). In cell monolayers expressing E-cadherin-GFP, we observed aligned trajectories of neighboring cells. Cells expressing only the SNAP-E-cadherin displayed more misaligned trajectories. Strikingly, the addition of the DNA linker restored the coordinated migration, demonstrating that the linkage by the DNA-E-cadherin hybrid system was translated to cell collectives (**Figure S5 A, Video 4**). We quantified the collective dynamics by calculating the velocity correlation length for the different conditions (**Figure S5 B, C**). It is a measure of coordinated motion inferred from vector fields mapped by particle image velocimetry (Petitjean et al., 2010) (**Figure 4 D**). The migration was analyzed for 7 hours, because the DNA linker was mainly internalized at this point (**Figure S5 E**). The expression of SNAP-E-cadherin resulted in a decreased correlation length of 135.8 ± 6.2 µm compared to cells expressing E-cadherin-GFP (159.4 ± 7.9 µm), in line with recent reports (Balasubramaniam et al., 2021; Das et al., 2015). The correlation length could be recovered completely through addition of the DNA linker (157.7 ± 10.9 µm, **Figure 4 E**). The increased coordination between migrating cells demonstrated a large-scale transduction of mechanical forces (Ladoux and Mège, 2017; Trepat and Sahai, 2018) facilitated by the DNA-E-cadherin hybrid system. To exploit the potential of DNA nanotechnology, we tested DNA linkers of different binding strengths (3.2, 10.4 and 17.7 kcal/mol). This revealed that the coordination between migrating cells directly depends on the molecular binding strength, since the correlation length increased from 138.3 ± 7.3 µm to up to 157.7 ± 10.9 µm when using stronger linkers (**Figure 4 F**). Furthermore, we functionalized the cellular membranes with the 17.7 kcal/mol linker using cholesterol-tags, which did not result in an increased correlation length (137.7 ± 8.9 µm **Figure 4 F**). This proves that the E-cadherin component of the hybrid system is crucial for its functionality.

In conclusion, we present a novel approach to investigate cell-cell adhesion by combining the advantages of DNA nanotechnology and protein engineering. The molecular binding strength of the DNA-E-cadherin hybrid is freely tunable by using different DNA sequences, but retains downstream signaling capabilities through the remaining domains of E-cadherin. Using single-cell force spectroscopy, we have demonstrated that the addition of the DNA linker leads to an increased cell-cell adhesion strength compared to the truncated E-cadherin. Furthermore, the simple addition of benzylguanine-tagged DNA to the culture medium provides reversibility and temporal control over cell-cell adhesion, since the linker strands can be removed again from the DNA-E-cadherin hybrid within seconds using strand displacement. Cell adhesion mediated by the DNA linker is fast and occurs within one hour. We show that the DNA linker system provides E-cadherin downstream signaling in terms of the formation of E-cadherin/β-catenin complexes, actin remodeling and mechanotransduction. Besides investigating cellular reactions, we demonstrate that our approach is suitable for studies on cell collectives, like monolayer migration.

Thus, the work presented here could be useful to directly assess the influence of cell-cell adhesion strength on tissue homeostasis (Guillot and Lecuit, 2013) and adhesion-dependent signaling (Charras and Yap, 2018). Since any DNA strand can be bound to SNAP-E-cadherin, our approach opens up the possibility to use DNA origami (Akbari et al., 2017) to link epithelial cells in a functional way. This would e.g. allow to investigate the influence of controlled E-cadherin spacing (Hellmeier et al., 2021) and therefore *cis-*clustering on adhesion and signaling processes. Moreover, the general concept of using a DNA-protein hybrid to tune molecular binding strengths could potentially be applied to investigate other binding-strength dependent cellular processes, e.g. cell-matrix adhesion through the modification of integrin (Kechagia et al., 2019). In summary, our approach based on the DNA-E-cadherin hybrid system allows to investigate (i) the immediate effect of a freely tunable molecular binding strength with temporal control and reversibility on different biological processes ranging from single cells to cell collectives, while (ii) maintaining the outside-in biochemical signaling activity of transmembrane receptors.

## Author contributions

A.S., K.G. and E.A.C.-A. conceived the project and designed the experiments. A.S. and N.H. performed the experiments and analyzed the results. C.L.G. acquired and analyzed the AFM data. D.O. designed the SNAP-E-cadherin plasmid and generated the cell lines. K.J. acquired STED images. A.S., K.G. and E.A.C.-A wrote the manuscript with input from all authors.

## Acknowledgements

We thank the members of the Biophysical Engineering Group and the Growth Factor Mechanobiology Group at the MPI for Medical Research for helpful discussion. We thank Jennifer Stow and René-Marc Mège for providing materials, Raimund Jung for help with cell sorting and the Optical Microscopy Facility at the MPI for Medical Research. K.J. thanks the Carl Zeiss Foundation for financial support. K.G. received funding from the Deutsche Forschungsgemeinschaft (DFG; German Research Foundation) under Germany’s Excellence Strategy via the Excellence Cluster 3D Matter Made to Order (EXC-2082/1-390761711) and the Max Planck Society. E.A.C.-A. acknowledges support from the DFG (SFB1129 P15) and the Baden-Württemberg Stiftung (3D MOSAIC). The Max Planck Society is appreciated for its general support.

## Notes

The authors declare no competing financial interest.

