## Supporting information for "Tuning epithelial cell-cell adhesion and collective dynamics with functional DNA-E-cadherin hybrid linkers"

### Materials and Methods

Design of the DNA linkers

All DNA linkers were designed and their hybridization strength was calculated using the Nucleic Acid Package (NUPACK) (Zadeh et al., 2011). It provides a thermodynamic analysis of interacting nucleic acid strands and calculates the free energy in kcal/mol of the secondary structure as well as equilibrium concentrations at a given temperature.

We designed our linker systems to consist of a 15-bp long anchor strand functionalized with either cholesterol (chol) or benzylguanine (bg). To the anchor strand, complementary, 30-bp-long linker strands (Strand 1 and Strand 2) with different hybridization strengths were bound. For the initial design of the 17.7 kcal/mol linker (called DNA linker in the manuscript until Figure 4), the *Design* function in NUPACK was used to generate ideal DNA sequences without unwanted self-complementarity.

The hybridization sequence for the 3.2 kcal/mol linker was manually generated by using 11 adenine/thymine bases. The 10.4 kcal/mol linker consists of 13 adenine/thymine bases. The sequence for toehold mediated strand displacement was described elsewhere (Zhang and Winfree, 2009). This toehold sequence was added to the sequence of the DNA linker 1. The invader strand is the complementary strand to the DNA linker 1 with the toehold overhang (called Reversible DNA linker 1).

<u>DNA sequences</u>

All custom DNA strands were purchased from Biomers (Ulm, Germany, purification: HPLC). Some of the linker strands were modified with a fluorophore. All DNA strands were diluted in Milli-Q water (Merck, Germany) to a stock concentration of 100 µM and stored at -20C until use.

*Anchor strand:*

5'/GTTCACAAGAAAGCG/3’/Chol

5'/GTTCACAAGAAAGCG/3’/BG

*17.7 kcal/mol linker (called DNA linker) strand 1:*

5’/CGCTTTCTTGTGAACACTCTTTCACTATCT/3'

*17.7k kcal/mol linker (called DNA linker) strand 2:*

Cy5/5‘/CGCTTTCTTGTGAACAGATAGTGAAAGAGA/3'

*10.4 kcal/mol linker strand 1:*

5‘/CGCTTTCTTGTGAACTTTTTTTTTTTTT/3'

*10.4 kcal/mol linker strand 2:*

5'/CGCTTTCTTGTGAACAAAAAAAAAAAAA/3'

*3.2 kcal/mol linker strand 1:*

5'/CGCTTTCTTGTGAACTTTTTTTTTTT/3'

*3.2 kcal/mol linker strand 2:*

5'/CGCTTTCTTGTGAACAAAAAAAAAAA/3'

*Reversible DNA linker strand 1:*
Atto647N/5'/CGCTTTCTTGTGAACACTCTTTCACTATCTTCTCCATGTCACTTC/3'

*Invader strand:*
5'/GAAGTGACATGGAGAAGATAGTGAAAGAGTGTTCACAAGAAAGCG/3'

<u>DNA linker assembly</u>

Duplexes of anchor and linker strand were pre-annealed by mixing 5 µl anchor strand, 5 µl linker strand, 20 µl $MgCl_2$ (100 mM, catalog no. M8226, Sigma-Aldrich) and 20 µl phosphate-buffered saline (PBS). This resulted in a final DNA concentration of 10 µM per strand. For duplex formation, the mix was put in a thermocycler (Bio-Rad), heated to 65 °C for 5 min and cooled down stepwise (10 °C every 30 sec) to 5 °C.

<u>Plasmids and cloning</u>

The plasmid coding for full-length E-cadherin-GFP was kindly provided by Jennifer Stow (Institute of Molecular Biosciences, University of Queensland, Addgene plasmid #28009). The plasmid coding for SNAP-E-cadherin345-mCherry, a truncated E-cadherin with an intracellular mCherry tag, was created via Gibson Assembly (GA). A SNAP-tag was introduced in the E-cadherin sequence between R154 and N376, replacing the extracellular domains EC1 and EC2 while maintaining the ER-import sequence and furin cleavage site of the E-cadherin prodomain.
In brief, the plasmid backbone including the E-cadherin prodomain was amplified from the template plasmid E-cadherin-EGFP-Halo (pDO33, unpublished data) via PCR using primers Fwd: *GAA TTC TAG AGG GCC CTA TTC TAT AGT GTC ACC TAA ATG CTA GAG CTC GC* and Rev: *TCT CTT CTG TCT TCT GAG GCC AGG AGA GGA GTT GGG AAA TGT GAG C*. From the same template the truncated E-cadherin sequence was amplified using primers Fwd: *ATC CGC GTT TAA ACT CGA GGT TAA TAA TCC CAC CAC GTA CAA GGG TCA GG* and Rev: *CCT TGC TCA CCA TAC TTC CTC CTC CTC CGT CGT CCT CGC CGC CTC CG.* The SNAP and mCherry fragments were amplified from template plasmid SNAPf-mCherry (pDO13,

unpublished data) using primers Fwd: *TGG CCT CAG AAG ACA GAA GAG AGA CAA AGA CTG CGA AAT GAA GC* and Rev: *CCT TGT ACG TGG TGG GAT TAT TAA CCT CGA GTT TAA ACG C* for SNAP and Fwd: *CGGA GGC GGC GAG GAC GAC GGA GGA GGA GGA AGT ATG GTG AGC AAG G* and Rev: *TGA CAC TAT AGA ATA GGG CCC TCT AGA ATT CTT ACT TGT ACA GCT CGT CCA TGC C*. All four fragments were assembled using the 2 x GA Master Mix (catalog no. E2611, NEB) following the manufacturer's instructions.

<u>Cell culture</u>

Jurkat T-cells (catalog no. TIB-152, ATCC) were grown in culture medium (RPMI, catalog no. 11875093, Gibco) supplemented with 10 % Fetal Bovine Serum (FBS, catalog no. 11140035, Gibco) and 1% Penicillin-Streptomycin (catalog no. P4333 Sigma Aldrich) at 37°C and 5% $CO_2$. They were split every 2-3 days. Before further processing, cells were centrifuged (5 min, 750 rpm), resuspended in phosphate-buffered saline (PBS, catalog no. 10010023, Gibco), centrifuged and finally resuspended in culture medium.
A431D cells initially described by (Lewis et al., 1997) were kindly provided by the group of René-Marc Mège (Institut Jacques Monod, Université de Paris). They were grown in culture medium (DMEM, catalog no. 1188002, Gibco) supplemented with 10% Fetal Bovine Serum (FBS, catalog no. 10270106, Gibco) and 1% Penicillin-Streptomycin (catalog no. P4333, Sigma Aldrich) at 37 °C and 5% $CO_2$. They were passaged every 2-3 days using 0.05% Trypsin (catalog no. 9002077, Merck). Before further processing, the culture medium was aspirated and cells were rinsed with PBS to remove dead cells and debris.

<u>Generation of SNAP-E-cadherin-mCherry and E-cadherin-GFP cell lines</u>

A431D cells have been transfected with either E-cadherin-GFP or SNAP-E-cadherin-mCherry via electroporation using an Amaxa Nucleofector I Device (Lonza, Cologne) with the Amaxa Cell line Nucleofector Kit T (program X-01; Lonza) according to the manufacturers protocol. Starting one day after transfection, cells were incubated in antibiotic selection medium (DMEM supplemented with 10 % FBS and 750 µg/ml geneticin (catalog no. 10131035, Gibco) for two weeks. Finally, antibiotic selected cells were sorted via fluorescence activated cell sorting (FACS) using a BD FACS Melody with 3 lasers (488/561/640) and 8 colors (2-2-4) configuration.

Linking of cells

Cells were incubated with linker medium (cultivation medium containing 10 mM $MgCl_2$ and 1 µM pre-annealed linker dsDNA) for 1 h at standard conditions (37 °C, 5% CO2) for all experiments except the force spectroscopy and the subcellular localization of YAP. The pre-annealed linker dsDNA consists of an anchor strand functionalized with either cholesterol or benzylguanine bound to Linker strand 1 or Linker strand 2. Fixed samples were mounted on a laser scanning confocal microscope (Zeiss LSM 900, Oberkochen, Germany) equipped with a 63x oil objective (Plan-Apochromat 63×/1.4 Oil DIC M27) and an Airyscan 2 module. Z-stacks of images were taken using the Airyscan mode with as step size of 0.13 µm. Within the ZEN software, an automated deconvolution was performed on the Airyscan data. All images were visualized using Fiji (Schindelin et al., 2012). For the visualization, maximum projections of the z-stacks were generated and brightness and contrast were adjusted. Live-cell timelapse videos were acquired using a laser scanning confocal microscope (Zeiss LSM 880, Oberkochen, Germany) equipped with a 20x air objective (LD A-Plan) and temperature and $CO_2$ control.

Sample preparation for single-cell force spectroscopy

2-well culture inlets (catalog no. 81176, Ibidi) were placed in a glass bottom atomic force microscopy dish (catalog no. GWST-3512, WillCo Wells). 2 000 cells (SNAP-E-cadherin or E-cadherin-GFP) were seeded into each inlet and incubated overnight at standard culture conditions. To prevent cell adhesion, the area outside the inlet was coated with 0.1 µg/ml Poly-L-lysine-Poly-ethylene-glycol (PLL(20)-g[3.5]-PEG(2), SuSoS, Switzerland). On the next day, the area outside the inlet was washed with PBS and dried at 37 °C. If no DNA linker was used for the experiment, adherent cells within the inlet were washed with warm PBS. The inlet was removed and the whole dish was filled with 3 ml medium. Freshly trypsinized cells were added to the dish excluding the area occupied by adherent cells.

When using the DNA linker, the adherent cells within the inlet were washed with warm PBS, the cultivation medium was replaced with linker medium containing 1 µM DNA Linker strand 1 and incubated for 1 h at standard conditions. In parallel, freshly trypsinized cells were incubated with linker medium containing 1 µM DNA Linker strand 2. The adherent cells were washed with warm PBS. The inlet was removed and the whole dish was filled with 3 ml culture medium. The trypsinized cells were added to the dish excluding the area occupied by adherent cells.

Single-cell force spectroscopy

For single-cell force spectroscopy, we used a NanoWizard 3 (JPK instruments, Bruker, Berlin), equipped with a CellHesion module (JPK instruments, Bruker, Berlin) and temperature and $CO_2$ control. Tipless cantilevers (catalog no. MLCT-O10, Bruker, Berlin) were functionalized with concanavalin-A (ConA-biotin; catalog no. C2272, Sigma) to facilitate cell capturing, adapting the protocol from (Friedrichs et al., 2010). Briefly, the cantilevers were cleaned for 15 min in an ultraviolet radiation and ozone (UV-O) cleaner (Jetlight) and placed in a Petri dish covered with parafilm. The cantilevers were then incubated overnight in 50 µL droplets of biotin-BSA (1 mg $ml^{-1}$ in $NaHCO_3$ buffer, 100 mM, pH 8.6, catalog no. A6043, Sigma), after which they were washed three times by immersion in fresh PBS. 50 µL droplets of streptavidin (1 mg $ml^{-1}$ in PBS, catalog no. S4762, Sigma) were added to the cantilevers. After 30 min incubation, the cantilevers were washed three times in PBS and placed in 50 µL droplets of ConA-biotin for additional 30 min. Then, the cantilevers were washed three times with PBS and placed in fresh PBS until use.

Immediately before use, cantilevers were calibrated with the thermal tune method (Schäffer, 2005) ($k = 0.01 - 0.03$ N $m^{-1}$). The experiment was performed in standard culture medium. For catching cells, the cantilever was brought into contact with a non-adherent cell at 1-3 nN constant force for 60-120 sec, after which the cell-functionalized cantilever was retracted and allowed to recover for 10 min before measuring cell-cell interactions. Then, the cell-functionalized cantilever was brought in contact with an adherent cell at 1 nN constant force for 2, 5 or 10 sec, before retraction, leading to the acquisition of a force-distance cycle. For each cell-functionalized cantilever, the process was repeated with 5 - 10 adherent cells. Between every measurement, cells were left to recover for 1 min. Detachment forces were calculated using the JPKSPM

Data Processing software. The values were plotted using GraphPad Prism 9. Error bars show the standard deviation. Plots generated from 3 independent experiments ($N$ = 3). Number of measured cells: $n$(E-cadherin-GFP) = 28. $n$(SNAP-E-cadherin) = 37. $n$(SNAP-E-cadherin + DNA linker) = 29.

<u>Toehold-mediated strand displacement</u>

A431D-SNAP-E-cadherin cells were seeded on a glass bottom imaging dish (catalog no. 81218, Ibidi) to reach a confluency of about 90%. Before the experiment, they were rinsed with warm PBS. They were incubated with the Reversible linker strand 1 tagged with Atto647N and the complementary DNA linker strand 2 anchored with Benzylguanine for 1 h at 37 °C and 5% $CO_2$ in the presence of 10 mM $MgCl_2$. After washing the sample with warm PBS and replacing the linking medium with cultivation medium supplemented with 10 mM $MgCl_2$, it was transferred to a laser scanning confocal microscope (Zeiss LSM 900, Oberkochen, Germany) equipped with a 63x oil objective (Plan-Apochromat 63×/1.4 Oil DIC M27).
Time-lapse videos (every 30 sec) of the linker tagged with Atto657N and SNAP-E-cadherin-mCherry were acquired at for 2 min. Then, the invader strand was added to achieve a concentration of 10 μM (10x excess). The sample was imaged for additional 8 min. 8-bit images with a color depth of 255 intensity values were acquired.
Using Fiji (Schindelin et al., 2012), we defined the background as an intensity value of five and all pixels below this threshold in all images of the time-lapse were set to zero. Then, the mean grey value was calculated for every image in the time-lapse.
The intensity values were normalized to the fraction of the maximum intensity. The intensity mean of multiple positions was generated and plotted over time using GraphPad Prism 9. The error bars show the standard deviation of $N$ = 3 experiments and $n$ = 31 measurements.

<u>Live imaging of cell-cell adhesion formation</u>

A431D-SNAP-E-cadherin cells were seeded on a glass bottom imaging dish (catalog no. 81218, Ibidi) to reach a confluency of about 40%. Before the experiment, they were rinsed with warm PBS. To label the actin cytoskeleton, SiR actin and verapamil

(catalog no. SC001, Spirochome, diluted 1:1000) were added to the cultivation medium and cells were incubated for 1 h at 37 °C and 5% $CO_2$.
The sample was mounted on an epifluorescence microscope (DeltaVision Imaging System on Olympus IX71 inverted microscope) equipped with temperature and $CO_2$ control and a 60x oil objective (Olympus, Plan Apo, NA = 1.4). E-cadherin-SNAP-mCherry and SiR actin were visualized by using the appropriate filter sets. Furthermore, the bottom of the glass coverslip was imaged by Interference Reflection Microscopy (IRM) by using TRITC excitation and FITC emission. Time-lapse videos of cells without and with the linker (every 10 minutes) were taken at different positions of the dish for multiple hours with constant 37 °C and 5% $CO_2$. Image acquisition started 20 min after linker addition.

Staining of E-cadherin, actin and β-catenin

Cells were seeded on clean glass coverslips to reach a confluency of about 50 – 70% on the day of the experiment. They were rinsed with warm PBS and (if applicable) incubated for 1 h at 37 °C and 5% $CO_2$ with linker medium. Before fixation, the cells were rinsed twice with warm PBS. Fixation was carried out in 4% paraformaldehyde (PFA) for 10 min at room temperature. Cells were permeabilized in 0.4% Triton X-100 in PBS for 5 min followed by 3 x 5 min washing in PBS. To stain the actin cytoskeleton, samples were incubated with Phalloidin-coumarin (catalog no. P2495, Sigma) diluted 1:200 in 5% BSA in PBS for 1h at room temperature.
For indirect immunostaining, samples were blocked with 5% BSA in PBS for 1 h at room temperature. Binding of primary antibodies was achieved by incubating the samples as it follows:
Anti-β-catenin mouse antibody (catalog no. 610153, BD) was diluted 1:100 in 5% BSA in PBS and incubated for 1 h at room temperature. Anti-E-cadherin mouse antibody (catalog no. sc-8426, Santa Cruz for E-cadherin-GFP, catalog no. 610181, BD for SNAP-E-cadherin-mCherry) was diluted 1:100 in 5% BSA in PBS and incubated for 1 h at room temperature. The samples were washed 3 times 5 min with PBS and incubated with anti-mouse donkey antibody conjugated to AF647 (catalog no. A-31571, Thermo Fisher), diluted 1:200 in 5% BSA in PBS and incubate for 1 h at room temperature. Subsequently, samples were washed 3 times 5 min with PBS. To

preserve fluorescence, samples were mounted with Mowiol 488 (catalog no. 81381, Sigma).

STED microscopy

Fixed cells stained for E-cadherin-GFP or SNAP-E-cadherin with an antibody conjugated to AF647 were imaged on an Abberior expert line microscope (Abberior Instruments GmbH, Germany) with a pulsed STED line at 775 nm using an excitation laser at 640 nm and spectral detection. The detection window was set to 650-725 nm to detect AF647-conjugated antibodies. Images were acquired with a 100×/1.4 NA magnification oil immersion lens (Olympus). The pixel size was set to 20 nm and the pinhole was set to 1 AU. The confocal as well as the STED laser power were set to 10%. For visualization, the contrast was adjusted using Fiji.

Colocalization of E-cadherin and β-catenin

Fixed samples were mounted on a laser scanning confocal microscope (Zeiss LSM 900, Oberkochen, Germany) equipped with a 63x oil objective (Plan-Apochromat 63×/1.4 Oil DIC M27) and an Airyscan 2 module). For the visualization of F-actin and E-cadherin, a background subtraction using a 100-pixel sliding paraboloid was used. The colocalization of β-catenin and E-cadherin was quantified by acquiring line plot profiles (length = 5 µm, width = 20 pixel) at the same positions in both channels in a single z-slice. The lines were manually placed in a way that the maximum intensity was at the middle. The intensity distribution was normalized to the fraction of the maximum value. Multiple line plots were averaged and plotted as intensity over distance using GraphPad Prism 9. The error bars show the standard deviation of $N$ = 3 preparations and $n$(E-cadherin-GFP) = 20, $n$(SNAP-E-cadherin-mCherry) = 22 and $n$(SNAP-E-cadherin-mCherry + DNA linker) = 23 measurements.

Subcellular localization of YAP

To assess the localization of Yes associated protein (YAP), cells were seeded on clean glass coverslips to reach a confluency of about 95 - 100% on the day of the experiment. The cells were cultured, incubated with DNA linkers for 3 h, fixed, permeabilized and blocked as described in the section above (Staining of actin and β-catenin). Samples were incubated with Anti-YAP mouse antibody (1:100, catalog no. 101199, Santa Cruz Biotechnology) for 1 h in 5% BSA in PBS and washed 3 times 5 min with PBS. Subsequently, 1 h incubation with anti-mouse donkey antibody conjugated to AF647 (catalog no. A-31571, Thermo Fisher) diluted 1:200 in 5% BSA in PBS was performed. After washing (3 times 5 min) with PBS, cells were counter-stained with DAPI (catalog no. D1306, Thermo Fisher) diluted 1:2000 in PBS for 15 min. Afterwards, samples were washed 3 times 5 min with PBS. To preserve fluorescence, samples were mounted with Mowiol 488 (catalog no. 81381, Sigma).

Images of 25 – 30 cells per field of view were acquired at a laser scanning confocal microscope using a 63x oil objective (Zeiss LSM 900, Oberkochen, Germany) and visualized using Fiji. Subcellular YAP intensities were quantified using the ImageJ Macro *Intensity Ratio Nuclei Cytoplasm Tool* (RRID:SCR_018573). First, a background correction was performed with all parameters set to zero. The nuclei were segmented based on the DAPI staining. All areas not segmented as nuclei were defined as cytosolic. The cytosolic and the nuclear YAP fractions within the image were quantified and the ratios were calculated. Further data processing was performed in RStudio running R Version 4.0.3. Measurements with a nuclear/cytosolic ratio below 0.85 were classified as cytosolic, while a ratio above 1.15 indicated nuclear localization. All measurements in between were classified as uniform. Percentages of the three localization classes were calculated for all conditions (E-cadherin-GFP, SNAP-E-cadherin, 3.2 kcal/mol linker, 10.4 kcal/mol linker, 17.7 kcal/mol linker) and plotted using GraphPad Prism 9.

Collective migration experiments

2-well culture inlets (catalog no. 81176, Ibidi) were placed in glass bottom culture dishes. $5x10^4$ cells (A431D cells expressing SNAP-E-cadherin-mCherry or E-cadherin-GFP) were seeded in 80 µl culture medium into the inserts and incubated overnight at standard conditions. The cells were washed with warm PBS and the medium was replaced with medium containing the following DNA linkers: 3.2 kcal/mol, 10.4 kcal/mol and 17.7 kcal/mol linkers anchored with benzylguanine to SNAP-E-cadherin or the 17.7 kcal/mol linker anchored with cholesterol into the cell membrane. In the case where no linker was used, fresh culture medium was added. The cells were incubated

for 1 h at standard conditions. Then the inlet was removed, the cells were carefully rinsed with PBS and the linker medium was replaced with standard cultivation medium containing 10 mM $MgCl_2$.
The cells were transferred to a live cell epifluorescence microscope (Leica DMi8 (Wetzlar, Germany) equipped with a 10x phase-contrast air objective and an incubation chamber) and incubated at 37 °C and 5% $CO_2$. Time-lapse videos (every 10 minutes) of the free edge were taken at different positions of the sample for multiple hours. Trajectories of single cells were manually tracked using the Fiji plugin *Manual Tracking*.

Particle image velocimetry analysis and velocity correlation length calculation

Time-lapse videos were cropped to a ROI of 385 x 1332 µm and background subtraction was performed. Therefore, the minimal intensity of the stack was projected and then subtracted using the *Calculator Plus* plugin. The videos were rotated in a way that the cells migrate from the left to the right.
The individual, cropped and background-subtracted images of the stack were loaded into *JPIV* (https://eguvep.github.io/jpiv/index.html) run in a Python environment. Since A431D cells do not migrate fast, we performed particle image velocimetry analysis on images taken 1 h apart (compare image 1 with image 6, image 2 with image 7, etc.) using first a 64x64 and then a 32x32 pixel interrogation window. The vector was placed in the middle of the 32x32 window. This generated a vector field with 32x32 pixel-spaced vectors. The vectors fields were batch-filtered by performing a normalized median test and a median filter, where all invalid vectors were excluded. These invalid vectors were replaced by the median.
A custom-written Python script was used to format the JPIV data for further processing. Finally, the velocity of the cell sheet and the velocity correlation were calculated in MATLAB using a script described elsewhere (Das et al., 2015), which was modified for the analysis of time-lapse video microscopy (Ollech et al., 2020).
In brief, the displacement vectors were divided by the time difference between the two images from which they were generated, resulting in the velocity vector $\boldsymbol{r}_{i,j}$, which was assigned to the central coordinate (*i,j*) of each 32x32 window. Since the axial migration is the dominant migration direction in the described experimental setup, only the lateral

component $\boldsymbol{U}_{i,j}$, perpendicular to the migration direction was used to calculate the velocity fluctuations $\boldsymbol{u}_{i,j}$ as:

$$\boldsymbol{u}_{i,j} = \boldsymbol{U}_{i,j} - \sum_{i=1,m} \sum_{j=1,n} \frac{\boldsymbol{U}_{i,j}}{m \times n} = \boldsymbol{U}_{i,j} - \boldsymbol{U}_{mean}$$

$U_{mean}$ is the mean velocity along the migration front. The lateral correlation function $\boldsymbol{C}_r$ was calculated as:

$$\boldsymbol{C}_r = \frac{\langle \boldsymbol{u}(r') * \boldsymbol{u}(r' + r)\rangle_{r'}}{\sqrt{\langle \boldsymbol{u}(r')^2\rangle} * \langle \boldsymbol{u}(r' + r)^2\rangle}$$

⟨…⟩ is the average and $r = || \boldsymbol{r}_{i,j} ||$ is the norm of $\boldsymbol{r}_{i,j}$. The first crossing of the threshold 0.01 with the lateral correlation function $\boldsymbol{C}_r$ was defined as the velocity correlation length. The mean correlation length was calculated for each ROI from 20 consecutive individual measurements corresponding to images acquired between 3.5 and 7.3 h after removing the confinement.

<u>Statistical analysis</u>

In GraphPad Prism 9, a one-way ANOVA significance test with Welch's correction was performed on the means of the different experiments using Dunnett's multiple comparison test. Thereby, the threshold for significance was defined as $\alpha = 0.05$. p-values between 0.1 and 0.01 correspond to (*), p-values between 0.01 and 0.001 correspond to (**), p-values between 0.001 and 0.0001 correspond to (***) and p-values < 0.0001 correspond to (****). No statistical significance is denoted as (n.s.). The individual datapoints were plotted together with the results of the statistical analysis using GraphPad Prism 9.

*Supporting Figure 1:* **Cells linked by cholesterol-anchored DNA on their surface.**

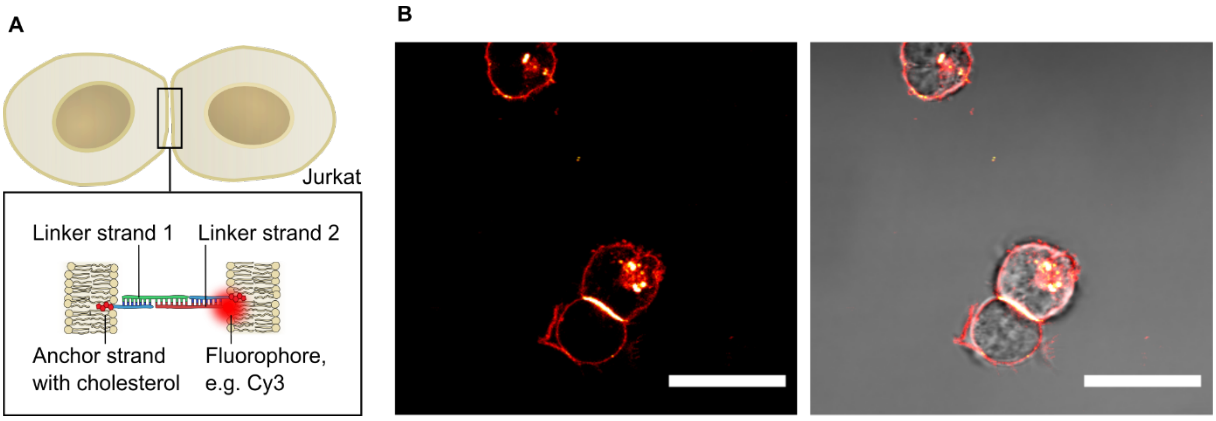


*Supporting Figure S1:* **Cells linked by cholesterol-anchored DNA on their surface. A** Sketch of linked Jurkat cells. Complementary linker strands are bound to an anchor strand which is functionalized with cholesterol. The linker strands can be tagged with a fluorophore, e.g. Cy3 for visualization. **B** Representative live-cell confocal and composite confocal and brightfield images of Jurkat cells incubated with the cholesterol-anchored DNA linker carrying Cy3 (*red*). Images representative of $N = 2$ independent experiments. Scale bars, 20 µm.

*Supporting Figure 2:* **Expression of SNAP-E-cadherin-mCherry and fluorescence decay over time.**

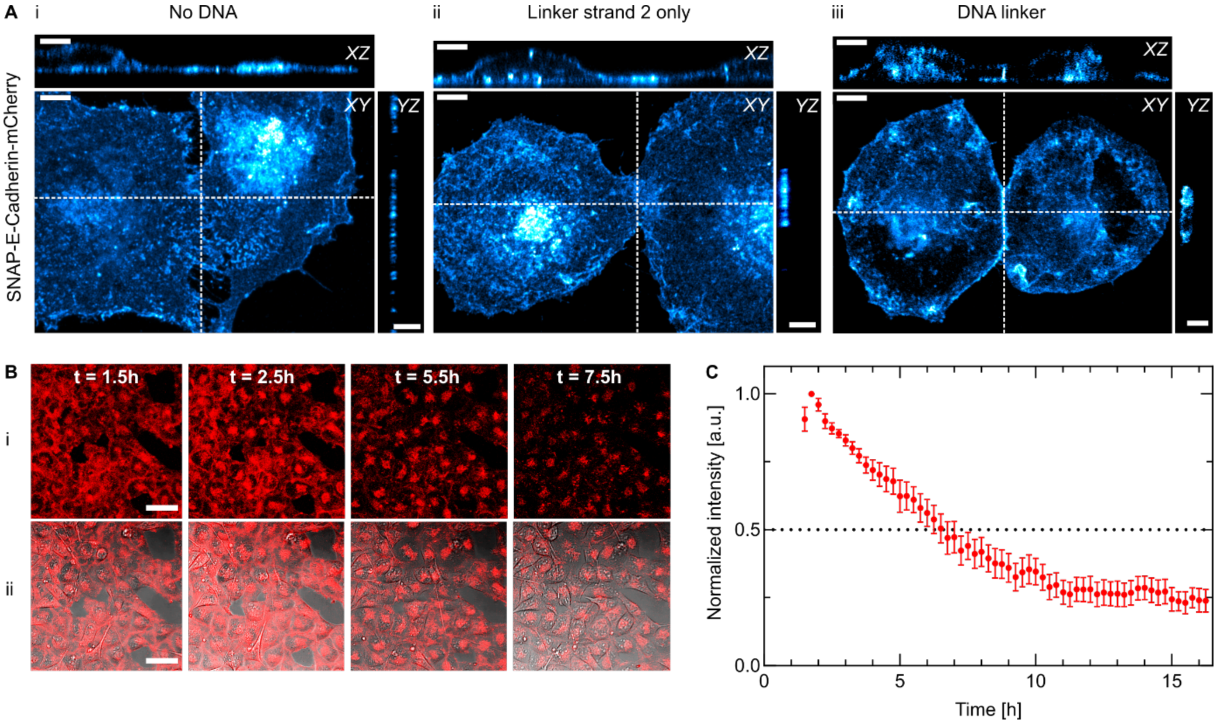


*Supporting Figure S2:* **A Expression of SNAP-E-cadherin-mCherry.** Whole-cell 3D reconstruction of the mCherry signal of A431D cells expressing SNAP-E-cadherin-mCherry (*cyan*). Cells were incubated in absence of DNA (**i**) with linker strand 2 only (**ii**) or the complete DNA linker (**iii**). (**ii**) and (**iii**) are the corresponding mCherry channels to **Figure 1 B**. Maximum projection and orthogonal slices through the positions indicated by the dashed line are shown. Scale bars, 5 µm. **B Fluorescence decay over time**. Selected timepoints of live-cell confocal (**i**) and composite brightfield and confocal (**ii**) images of SNAP-E-cadherin expressing cells pre-incubated with the Cy5 tagged DNA linker (*red*). Scale bars, 50 µm. **C** Quantification of the fluorescence decay over time starting at the beginning of image acquisition, 1.5 h after start of the incubation. Data obtained from one preparation (*N = 1*) and *n = 6* measurements. Error bars show the standard deviation.

.

*Supporting Figure 3:* **Single cell force spectroscopy**

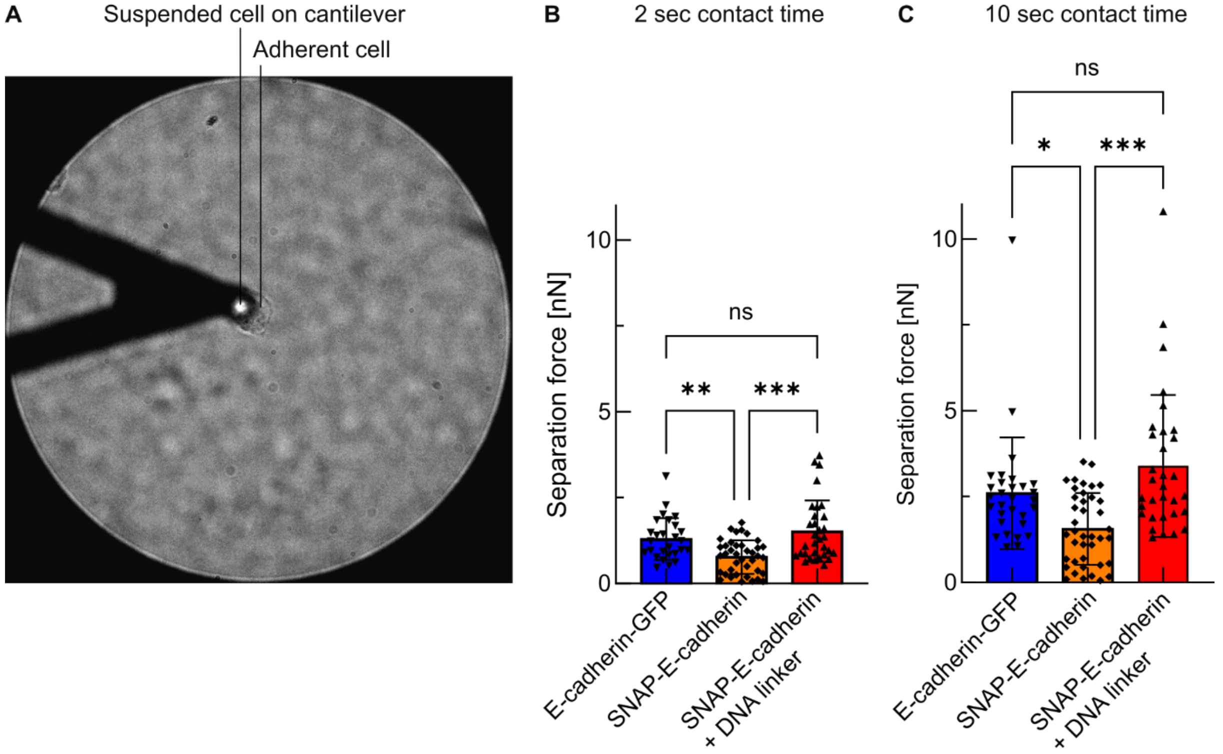


*Supporting Figure S3:* **Single cell force spectroscopy. A** Brightfield image showing a suspended cell bound to the AFM cantilever functionalized with Concanavalin A on top of an adherent cell. **B** Comparison of the separation forces for 2 seconds contact time. Bars show the mean (E-cadherin-GFP = 1.292 ± 0.607 nN; SNAP-E-cadherin = 0.772 ± 0.49 nN; SNAP-E-cadherin + DNA linker = 1.521 ± 0.894 nN). Error bars show the standard deviation. Plots generated from 3 independent experiments ($N$ = 3). Number of measured cells: $n$(E-cadherin-GFP) = 27. $n$(SNAP-E-cadherin) = 39. $n$(SNAP-E-cadherin + DNA linker) = 32. **C** Comparison of the separation forces for *10 seconds contact*. Bars show the mean (E-cadherin-GFP = 2.599 ± 1.619 nN; SNAP-E-cadherin = 1.562 ± 1.045 nN; SNAP-E-cadherin + DNA linker = 3.369 ± 2.065 nN). Error bars show the standard deviation. Plots generated from 3 independent experiments ($N$ = 3). Number of measured cells: $n$(E-cadherin-GFP) = 30. $n$(SNAP-E-cadherin) = 40. $n$(SNAP-E-cadherin + DNA linker) = 32. ns no significance. (*) p-value between 0.1 and 0.01. (**) p-value between 0.01 and 0.001. (***) p-value between 0.001 and 0.0001. Multiple ANOVA tests with Welch's correction. Alpha was set to 0.05.

409 *Supporting Figure 4:* **Super resolution images of E-cadherin at cell-cell contact.**

410

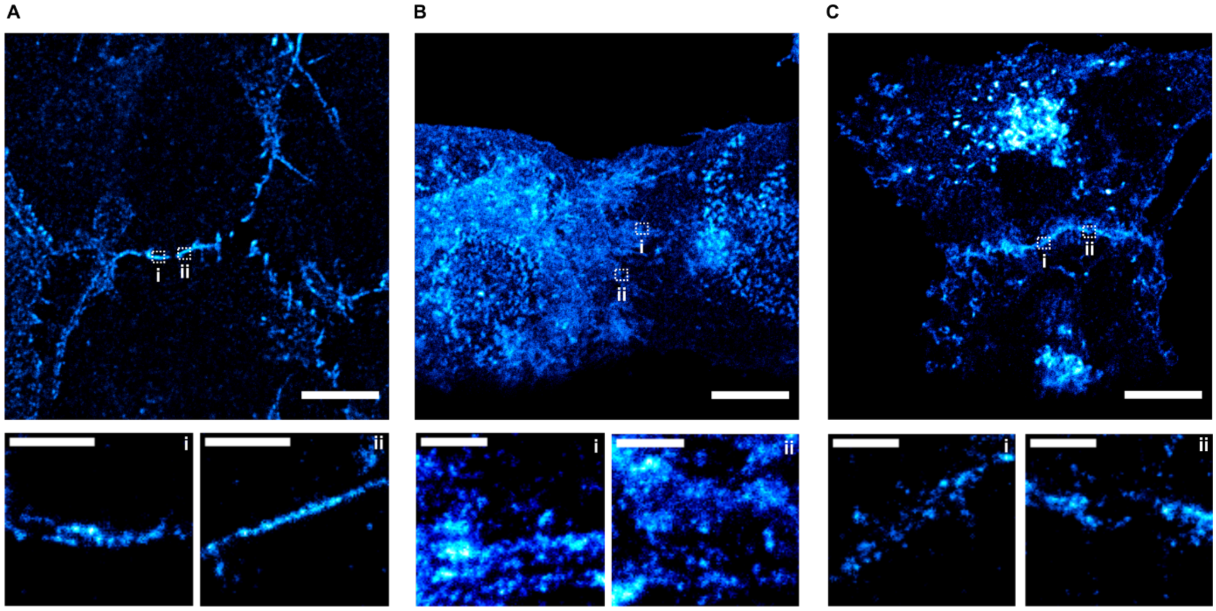


*Supporting Figure 4:* **Super resolution images of E-cadherin at cell-cell contact.** The intracellular domain of E-cadherin is visualized by indirect immunostaining with AF647. Top row: Overview confocal images, scale bar 10 µm. Bottom row: (**i**) and (**ii**) STED images of zoom-ins. The positions are indicated by the dashed lines in the overview image. Scale bar, 1 µm. **A** E-cadherin-GFP. **B** SNAP-E-cadherin. **C** SNAP-E-cadherin + DNA linker. Images representative of $N = 2$ independent experiments.

411

*Supporting Figure 5:* **Effect of the DNA-E-cadherin hybrid system on collective migration.**

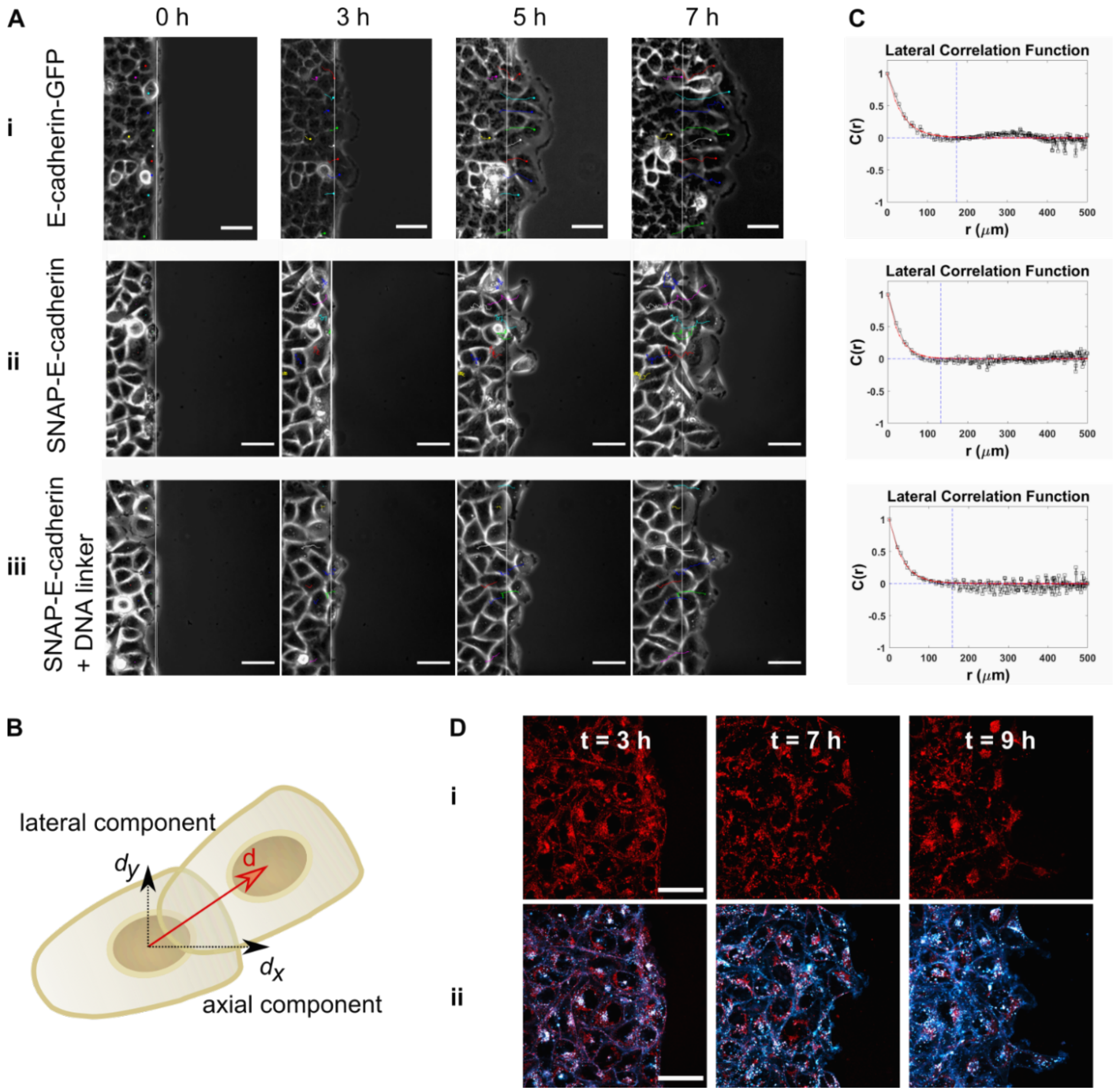


*Supporting Figure S5:* **Effect of the DNA-E-cadherin hybrid system on collective migration. A** A431D cells were cultured in confinement for collective migration assays. Live-cell time-lapse phase contrast images of the migration front taken at the indicated timepoints after removing the confinement. The dashed line shows the initial position of the collective front. Trajectories of single cells are shown by colored lines. Scale bars, 50 µm. (**i**) E-cadherin-GFP, (**ii**) SNAP-E-cadherin, (**iii**) SNAP-E-cadherin + DNA linker. **B** Principle of the particle image velocimetry (PIV) analysis: The velocity vector *d* and its axial ($d_x$) and lateral ($d_y$) component are calculated. Only the lateral component is used for the calculation of the correlation length. Adapted from *Ollech et. al., 2020*. **C** Representative plots of the lateral correlation function *C(r)* for (**i**) E-cadherin-GFP, (**ii**) SNAP-E-cadherin, (**iii**) SNAP-E-cadherin + DNA linker. The correlation length is defined as the distance *r* at the first zero-crossing of the function, as described previously by *Das et. al., 2015*. **D** Confocal images of the DNA linker (*red*, **i**) and SNAP-E-cadherin-mCherry (*blue;* **ii** overlay) at the migration front at the indicated timepoints. Scale bar, 50 µm.

### Video descriptions

*Supporting Video 1:* **3D projections of single cells.**
A431D cells expressing SNAP-E-cadherin-mCherry (*cyan*) were incubated with only linker strand 2 Cy5 (*red*) or the complete, Cy5-tagged DNA linker (*red*). Images correspond to Figure 1 B and Figure S2 A. The scale bar shows 5 µm.

*Supporting Video 2:* **Toehold-mediated strand displacement.**
A431D cells expressing SNAP-E-cadherin-mCherry (*cyan*) were incubated with a toehold-carrying DNA linker tagged Atto647N (*red*). The DNA linker diffuses out of focus after addition of the invader strand (added at t = 1.75 min). Video corresponding to Figure 2 B. The scale bar shows 50 µm.

*Supporting Video 3:* **Adherens junction formation.**
A431D cells expressing SNAP-E-cadherin (*cyan*) were incubated with the DNA linker. The video starts directly after the linker addition. Bottom of the cell (Interference Reflection Microscopy, IRM, *grey*) and actin stained with SiR actin (*red*) are shown. Video corresponding to Figure 2. The scale bar shows 10 µm.

*Supporting Video 4:* **Migration front of cell collectives.**
The collective migration of cells expressing SNAP-E-cadherin, without and with the DNA linker, was imaged in brightfield and individual cells were tracked manually. Video corresponding to Figure S5 A. The scale bar shows 50 µm.

### Methods References:


1. Das, T., Safferling, K., Rausch, S., Grabe, N., Boehm, H., and Spatz, J.P. (2015). A molecular mechanotransduction pathway regulates collective migration of epithelial cells. Nature Cell Biology *17*, 276-287.
2. Friedrichs, J., Helenius, J., and Muller, D.J. (2010). Quantifying cellular adhesion to extracellular matrix components by single-cell force spectroscopy. Nature Protocols *5*, 1353-1361.
3. Lewis, J.E., Wahl, J.K., III, Sass, K.M., Jensen, P.J., Johnson, K.R., and Wheelock, M.J. (1997). Cross-Talk between Adherens Junctions and Desmosomes Depends on Plakoglobin. Journal of Cell Biology *136*, 919-934.
4. Ollech, D., Pflästerer, T., Shellard, A., Zambarda, C., Spatz, J.P., Marcq, P., Mayor, R., Wombacher, R., and Cavalcanti-Adam, E.A. (2020). An optochemical tool for light-induced dissociation of adherens junctions to control mechanical coupling between cells. Nature Communications *11*, 472.
5. Schäffer, T.E. (2005). Calculation of thermal noise in an atomic force microscope with a finite optical spot size. Nanotechnology *16*, 664-670.
6. Schindelin, J., Arganda-Carreras, I., Frise, E., Kaynig, V., Longair, M., Pietzsch, T., Preibisch, S., Rueden, C., Saalfeld, S., Schmid, B., *et al.* (2012). Fiji: an open-source platform for biological-image analysis. Nature Methods *9*, 676-682.
7. Zadeh, J.N., Steenberg, C.D., Bois, J.S., Wolfe, B.R., Pierce, M.B., Khan, A.R., Dirks, R.M., and Pierce, N.A. (2011). NUPACK: Analysis and design of nucleic acid systems. Journal of Computational Chemistry *32*, 170-173.
8. Zhang, D.Y., and Winfree, E. (2009). Control of DNA Strand Displacement Kinetics Using Toehold Exchange. Journal of the American Chemical Society *131*, 17303-17314.
